# Mutations in efflux pump Rv1258c (Tap) cause resistance to pyrazinamide and other drugs in *M. tuberculosis*

**DOI:** 10.1101/249102

**Authors:** Jiayun Liu, Wanliang Shi, Shuo Zhang, Gail Cassell, Dmitry A. Maslov, Kirill V. Shur, Olga B. Bekker, Valery N. Danilenko, Xiaoke Hao, Ying Zhang

## Abstract

Although drug resistance in *M. tuberculosis* is mainly caused by mutations in drug activating enzymes or drug targets, there is increasing interest in possible role of efflux in causing drug resistance. Previously, efflux genes are shown upregulated upon drug exposure or implicated in drug resistance in overexpression studies, but the role of mutations in efflux pumps identified in clinical isolates in causing drug resistance is unknown. Here we investigated the role of mutations in efflux pump Rv1258c (Tap) from clinical isolates in causing drug resistance in *M. tuberculosis* by constructing point mutations V219A, S292L in Rv1258c in the chromosome of *M. tuberculosis* and assessed drug susceptibility of the constructed mutants. Interestingly, V219A, S292L point mutations caused clinically relevant drug resistance to pyrazinamide (PZA), isoniazid (INH), and streptomycin (SM), but not to other drugs in *M. tuberculosis*. While V219A point mutation conferred a low level resistance, the S292L mutation caused a higher level of resistance. Efflux inhibitor piperine inhibited INH and PZA resistance in the S292L mutant but not in the V219A mutant. S292L mutant had higher efflux activity for pyrazinoic acid (the active form of PZA) than the parent strain. We conclude that point mutations in the efflux pump Rv1258c in clinical isolates can confer clinically relevant drug resistance including PZA and could explain some previously unaccounted drug resistance in clinical strains. Future studies need to take efflux mutations into consideration for improved detection of drug resistance in *M. tuberculosis* and address their role in affecting treatment outcome in vivo.

## INTRODUCTION

Multidrug resistant (MDR) tuberculosis (MDR-TB), defined by resistance to at least two frontline drugs isoniazid and rifampin, poses a significant challenge to effective treatment and control of the disease. In 2015, there were at least 480,000 cases of MDR-TB and about 9.5% of MDR-TB cases were extensively drug resistant TB or XDR-TB (1). Drug resistance in the causative agent *M. tuberculosis* is mainly caused by mutations affecting enzymes involved in drug activation or by alterations or overexpression of drug targets (2). However, efflux pumps, which have been found to cause antibiotic resistance in many other bacteria (3) and also nontuberculous mycobacteria (4, 5) have only recently been shown to be involved in clinical drug resistance in *M. tuberculosis* in the case of clofazimine and bedaquiline (6, 7).

The efflux protein Rv1258c, also called Tap or P55, was previously shown to be involved in resistance to different drugs in overexpression studies involving *M. smegmatis* and *M. bovis* BCG (8, 9). For example, overexpression of Rv1258c was shown to confer resistance to INH, rifampin, ethambutol, PAS, ethambutol in *M. bovis* BCG (9). Additionally, Jiang et al. showed that Rv1258c was overexpressed upon exposure to isoniazid (INH) and rifampin (RIF) in MDR-TB clinical isolates and hypothesized that its overexpression may cause drug resistance in clinical TB strains (10). Furthermore, various single nucleotide polymorphisms (SNPs) in different efflux pump genes Rv0194, Rv1217, Rv1258, Rv1273, Rv1877, Rv1250, and Rv2688 in XDR-TB clinical isolates were recently identified (11). However, the relevance of these SNPs in causing drug resistance in clinical strains has been unclear. In this study, we identified SNPs in the efflux gene Rv1258c (tap) from clinical isolates of *M. tuberculosis* and then evaluated the significance of the SNPs in causing clinically relevant drug resistance by introducing these point mutations into the genome of the isogenic strain of *M. tuberculosis*. We demonstrate that mutations (V219A, S292L) in efflux pump Tap Rv1258c identified in clinical strains can cause clinically relevant drug resistance in *M. tuberculosis*.

## MATERIALS AND METHODS

### Bacterial strains, plasmids and drugs

*M. tuberculosis* H37Ra was grown at 37°C in Middlebrook 7H9 liquid medium or on 7H11 agar plates supplemented with 10% (v/v) albumin–dextrose–catalase (ADC, Becton Dickinson, Sparks, MD, USA) plus 0.5% (v/v) glycerol, 0.25% (v/v) Tween 80. Plasmids p2NIL and pGOAL19 used in this study were obtained from Addgene (Cambridge, MA, USA). Isoniazid (INH), rifampicin, streptomycin (SM), ethambutol, pyrazinamide (PZA), levofloxacin, amikacin, cycloserine, p-aminosalicylic acid (PAS), clofazimine (CFZ), tetracycline, linezolid, clarithromycin, and piperine were purchased from Sigma-Aldrich (St Louis, MO, USA). The drugs were dissolved in DMSO and further diluted to obtain the desired concentrations in culture media for drug susceptibility testing (see below).

### Construction of Rv1258c point mutation mutants by homologous recombination

*M. tuberculosis* mutants were constructed by homologous recombination as described previously (12). Briefly, the Rv1258c gene was amplified with its adjacent 1 kb fragment on both sides from *M. tuberculosis* genomic DNA and cloned into p2NIL plasmid vector. Then, the mutated constructs Rv1258c V219A, S292L were obtained by QuikChange II XL site-directed mutagenesis kit (Agilent, Santa Clara, CA, USA) with primers TPV219AF: 5’-TTCGCCTGGAACCTGCGGGTATTGCGCACCC-3’, TPV219AR: 5’-CCAGGCGAAGCGCAGCCCCTCGGCGATCCC-3’, TPS292LR: 5’-CCCCGTCGCGTGACCATGCTGACCGCGGTTCTTACCCTGGG-3’, TPS292LR: 5’-CCCAGGGTAAGAACCGCGGTCAGCATGGTCACGCGACGGGG-3’. pGOAL19 cassette was cloned into PacI site of p2NIL containing the mutant (V219A and S292L) version of the Rv1258c gene to form suicide delivery constructs. The suicide delivery plasmid DNA was subjected to 100 mJ/cm2 of UV irradiation followed by addition of 200 μl of electrocompetent mycobacteria. Electroporation was performed with the parameter 2.5 kV, 1000 Ω, 25 mF. The electroporated cells were added with 200 μl 7H9-Tween 80 recovery medium and incubated at 37°C for 24h followed by plating onto 7H11/ADC plates containing kanamycin at 20 μg/ml, hygromycin at 50 μg/ml, and X-Gal at 5μg/ml. The plates were incubated at 37°C for 4-5 weeks until colonies appeared. Single cross-over transformants were picked and restreaked onto fresh 7H11/ADC without any antibiotics and incubated at 37°C for 2-3 weeks. A loopful of bacteria from the plates were resuspended in 1 ml of 1-mm glass beads and 3 mL of 7H9/ADC/Tween 80 and vortexed vigorously. Serial dilutions were plated onto plates containing X-gal with and without sucrose at 2% (w/v) and incubated for 4 weeks until the potential double cross-over transformants appeared. The kanamycin-sensitive colonies were screened using colony-PCR followed by DNA sequencing to confirm the desired point mutation being constructed in the genome of *M. tuberculosis* H37Ra.

### Drug susceptibility testing and isoniazid/piperine combination studies

The Rv1258c V219A and S292L mutants and parent strain *M. tuberculosis* H37Ra were grown in7H9/ADC/Tween 80 medium at 37°C for 2-3 weeks (approximately 1×10^8^ colony-formingunits (CFU)/ml) when they were diluted in serial 10-fold dilutions and inoculated onto 7H11 agar plates containing different concentrations of drugs in triplicate. The following drug concentrations were used: SM (0.5, 1, 2, 4, 8 μg/ml). INH (0.125, 0.25, 0.5, 1, 2, 4, 8, 16, 32, 64 μg/ml), PZA (50, 100, 200, 400, 800, 1600 μg/ml, pH 6.0), p-aminosalicylic acid (PAS) (0.25, 0.5, 1, 2, 4, 8, 16, 32, 64, 128 μg/ml), rifampicin (0.25, 0.5, 1, 2, 4 μg/ml), ethambutol (1, 2, 4, 8, 16 μg/ml), levofloxacin (0.25, 0.5, 1, 2, 4 μg/ml), amikacin (0.5, 1, 2, 4, 8 μg/ml), cycloserine (5, 10, 20, 40, 80 μg/ml), clofazimine (CFZ) (0.125, 0.25, 0.5, 1, 2 μg/ml), tetracycline (0.5, 1, 2, 4, 8, 16, 32, 64, 128 μg/ml), linezolid (0.25, 0.5, 1, 2, 4 μg/ml), clarithromycin (0.125, 0.25, 0.5, 1, 2 μg/ml). The plates were incubated at 37°C in 5% CO_2_ for 3–4 weeks. The isoniazid/piperine combination study was performed on the Rv1258c V219A and S292L mutants with different concentrations of INH tested in the presence of increasing concentrations of piperine (0, 5, 10, 20, 40 μg/ml).

### PZA uptake and accumulation experiments

Two-week-old cultures of Rv1258c V219A and S292L mutants *M. tuberculosis* H37Ra, grown in in7H9/ADC/Tween 80 medium, were harvested and the cell pellets were resuspended in 7H9 medium (pH 6.6) at 5×10 cells/ml. The PZA uptake and accumulation study was performed as described previously (13). Briefly, [^14^C]PZA, purchased from Vitrax (Placentia, CA, USA), was added to the cell suspensions to a concentration of 1 μCi/ml and the cell mixtures were incubated at 37°C. At different time points, 50 μl portions were removed and washed with 1×PBS buffer (pH 6.6) with 0.1M LiCl by filtration on 0.45μm-pore-size nitrocellulose filters by using a vacuum pump. The amount of radioactivity associated with the bacterial cells was determined by autoradiography and scintillation counting.

## RESULTS AND DISCUSSION

### Identification of point mutations in Rv1258c from clinical isolates of *M. tuberculosis* and construction of Rv1258c point mutations in *M. tuberculosis*

To determine possible roles of efflux pumps in causing drug resistance, we performed database search for mutations in the efflux pump Rv1258c (Tap) among clinical isolates whose whole genome sequences are deposited in the PubMed NCBI database. A number of single nucleotide polymorphisms (SNPs) were found in the Tap efflux gene Rv1258c (D23V, V219A, S292L) in different clinical isolates of *M. tuberculosis*, which may affect the function of the efflux protein. This is consistent with the identification of Rv1258c P369T and G391R in XDR-TB clinical isolates from sequenced genomes of *M. tuberculosis* strains from Pakistan (11).

To address the role of the identified point mutations in Rv1258c, we attempted to construct the point mutations D23V, V219A and S292L by site-directed mutagenesis by PCR and the altered mutant sequences were confirmed to be correct by DNA sequencing. We were able to successfully construct Rv1258c point mutations V219A and S292L, but not D23V (for unknown reasons) in *M. tuberculosis* H37Ra. It is worth noting that the constructed Rv1258c point mutation V219A and S292L mutant strains had no growth defect or altered morphology as seen in Rv1258c deletion mutant strain (9). Therefore, we evaluated the susceptibility of Rv1258c point mutation V219A and S292L mutants to various drugs as described below.

### Rv1258c point mutations confer resistance to PZA, INH, and SM

The results of drug susceptibility testing showed that Rv1258c S292L mutant was more resistant to streptomycin, isoniazid, and pyrazinamide than the parent strain H37Ra (Table 1). However, Rv1258c V219A point mutation caused a lower level of resistance to the above drugs than the S292L mutation. It is noteworthy that while V219A mutation only caused a marginal level of INH resistance (MIC=0.125 μg/ml), S292L mutation caused a very high level of INH resistance (MIC=32 μg/ml) (Table 1). This latter finding is significant as it offers a possible alternative mechanism of INH resistance in addition to mutations in *katG* (2, 14) or *inhA (15*). Introduction of the V219A and S292L mutations in *M. tuberculosis* did not alter the susceptibility to other drugs including rifampicin, ethambutol, levofloxacin, amikacin, cycloserine, p-aminosalicylic acid (PAS), clofazimine (CFZ), tetracycline, linezolid and clarithromycin (Fig. 1-4, Table 1). Our finding that Rv1258c S292L point mutation conferred resistance to streptomycin and isoniazid is consistent with the previous observation of overexpression of Rv1258c (9). However, the Rv1258c S292L and V219A mutations did not alter susceptibility to other drugs such as tetracycline, RIF, EMB, CFZ, where previous studies have demonstrated that overexpression of Rv1258c could cause resistance to these drugs (5, 9, 16). One possibility for such a discrepancy is that the point mutations at S292L and V219A cause differential binding to different drugs such that they have differential effects on causing selective drug resistance to some but not other drugs as seen in overexpression study of the wild type functional protein Rv1258c (9, 16).

**TABLE 1.** Susceptibility of *M. tuberculosis* efflux pump (Tap, Rv1258c) point mutation mutants to different TB drugs

| Drugs | Drug susceptibility results# |  |
| --- | --- | --- |
|  | Rv1258c V219A | Rv1258c S292L |
| Isoniazid | R (MIC 0.125 µg/ml) | R (MIC 32 µg/ml) |
| Pyrazinamide | R (MIC 100 µg/ml) | R (MIC 800 µg/ml) |
| Streptomycin | S | R (MIC 4.0 µg/ml) |
| Rifampin | S | S |
| Ethambutol | S | S |
| Amikacin | S | S |
| Cycloserine | S | S |
| p-aminosalicylic acid (PAS) | S | S |
| Clofazimine | S | S |
| Tetracycline | S | S |
| Linezolid | S | S |
| Clarithromycin | S | S |

**Figure 1.**
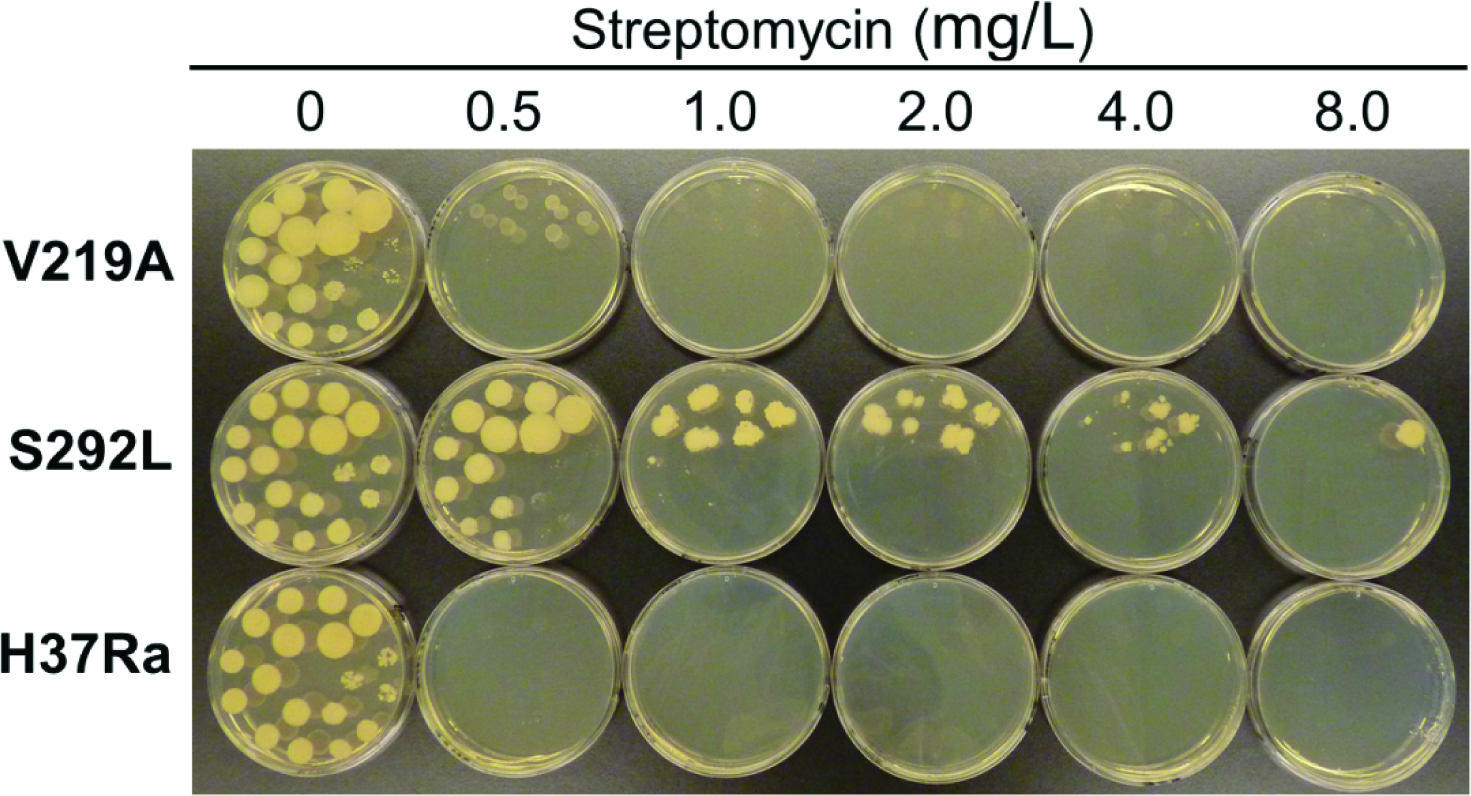
The Rv1258c S292L mutant has higher level of resistance to streptomycin than The V219 mutant.

**Figure 2.**
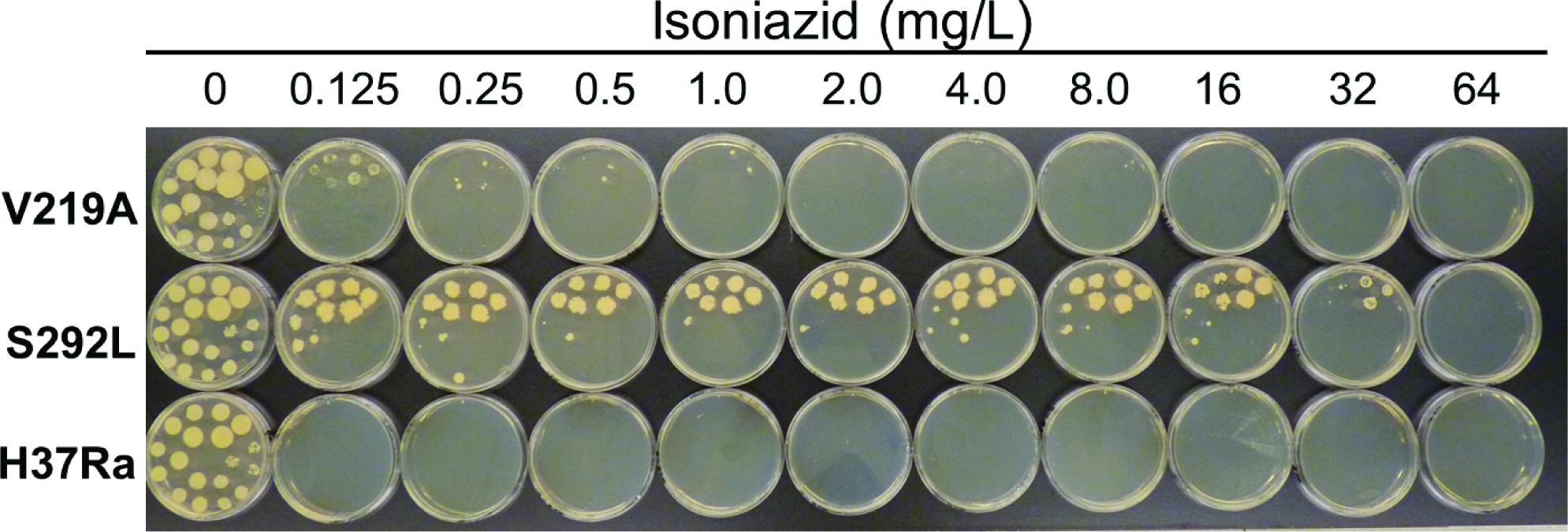
The Rv1258c S292L mutant is highly resistant to isoniazid, while the V219A mutant is only slightly resistant.

**Figure 3.**
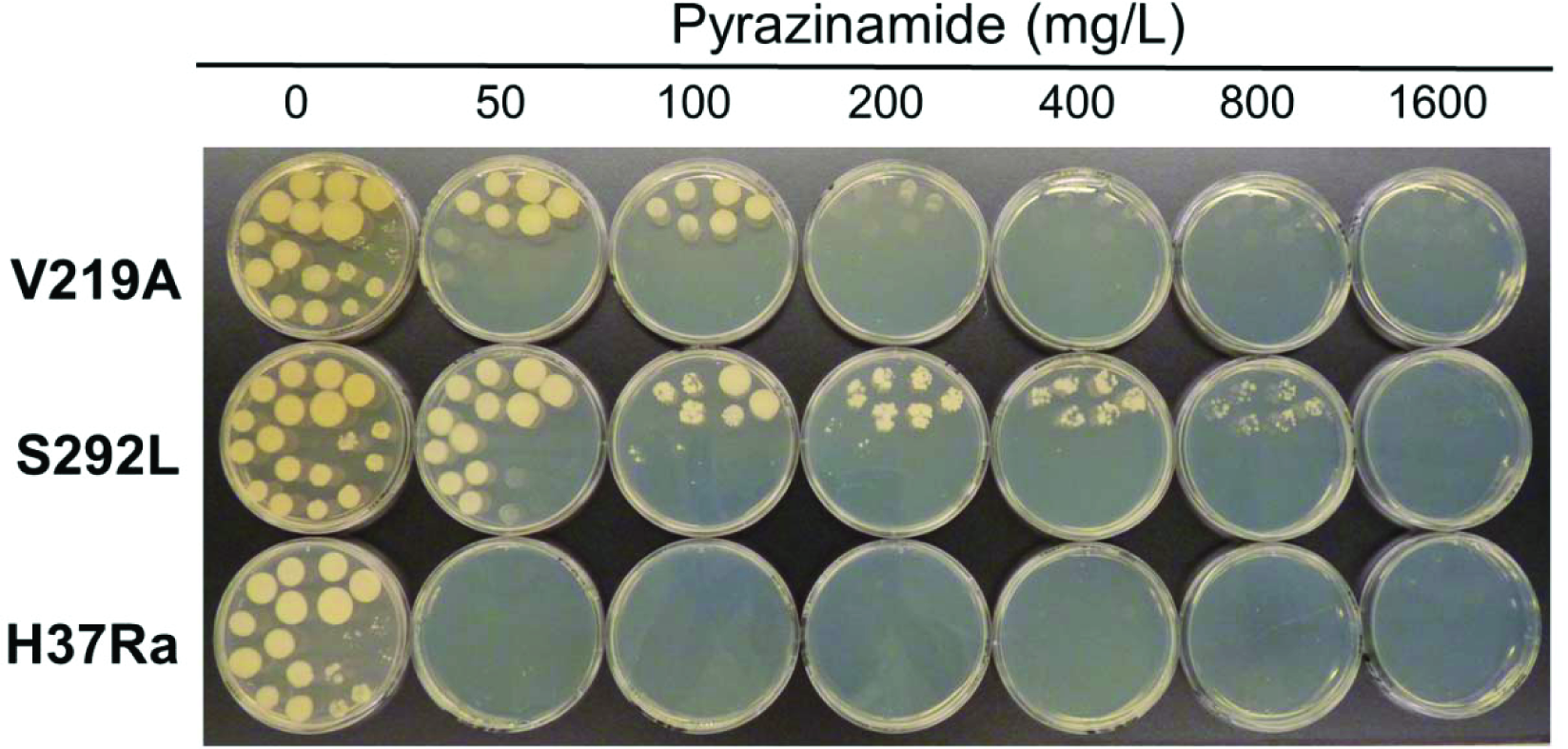
The Rv1258c S292L mutant has a higher level of PZA resistance than the V219A mutant.

**Figure 4.**
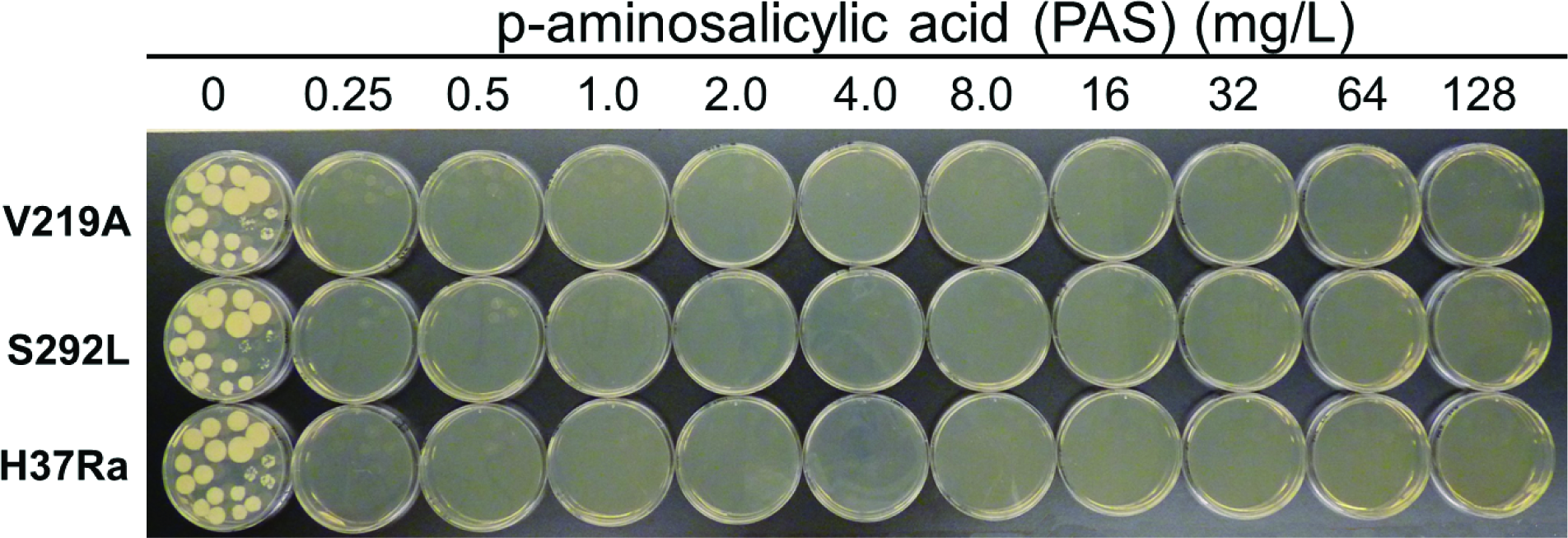
The Rv1258c S292L and V219A mutants remain susceptible to PAS.

Previous studies that evaluated drug resistance conferred by overexpression of Rv1258c did not test PZA, presumably because of the well-known problem with its susceptibility testing (17). Here, it is of interest to note that introducing the S292L point mutation in Rv1258c into the chromosome of *M. tuberculosis* conferred a high level of PZA resistance (800 μg/ml), whereas the V219A mutation caused a low level of PZA resistance (100-200 μg/ml) (Fig. 3). Although PZA resistance is mostly caused by *pncA* mutations (18) and less commonly by *rpsA (19), panD (20*), and *clpC1(21*, 22) mutations, some PZA-resistant *M. tuberculosis* strains without mutations in the above known genes do exist (23) (Zhang Y, unpublished). Recently, we have shown that overexpression of efflux proteins Rv0191, Rv3756c, Rv3008, and Rv1667c could all confer a low level PZA resistance in *M. tuberculosis (24*), indicating a role of efflux in PZA resistance. However, not all efflux pumps are involved in PZA resistance as overexpression of DrrAB in *M. tuberculosis* did not confer PZA resistance (Zhang Y, unpublished). Here it is worth noting that by constructing point mutations in the genome of the efflux pump gene Rv1258c, we were able to convincingly demonstrate that the S292L mutation is indeed causative of PZA resistance. This finding suggests that mutations in Rv1258c could be a potential new mechanism of PZA resistance in clinical isolates without known structural gene (*pncA, rpsA, panD*) mutations. Future studies are needed to determine how frequent such mechanism of PZA resistance mediated by mutations in Rv1258c occur in clinical isolates.

### Piperine inhibits the INH and PZA resistance in the Rv1258c S292L mutant

Piperine is a known inhibitor of Rv1258c in *M. tuberculosis* (25). To determine if the S292L point mutation mediated higher level of resistance is due to elevated efflux activity of the Rv1258c S292L mutant protein, we tested if piperine could antagonize the INH and PZA resistance mediated by the Rv1258c S292L mutant protein. The piperine/INH or piperine/PZA combination study showed that piperine reduced both INH and PZA resistance or increased INH and PZA susceptibility in the Rv1258c S292L mutant *M. tuberculosis* strain, but not in the V219A mutant strain (Fig. 5). The results indicate that the higher level of INH or PZA resistance in the Rv1258c S292L mutant is caused by higher efflux activity of the mutant protein that could be inhibited by piperine.

**Figure 5.**
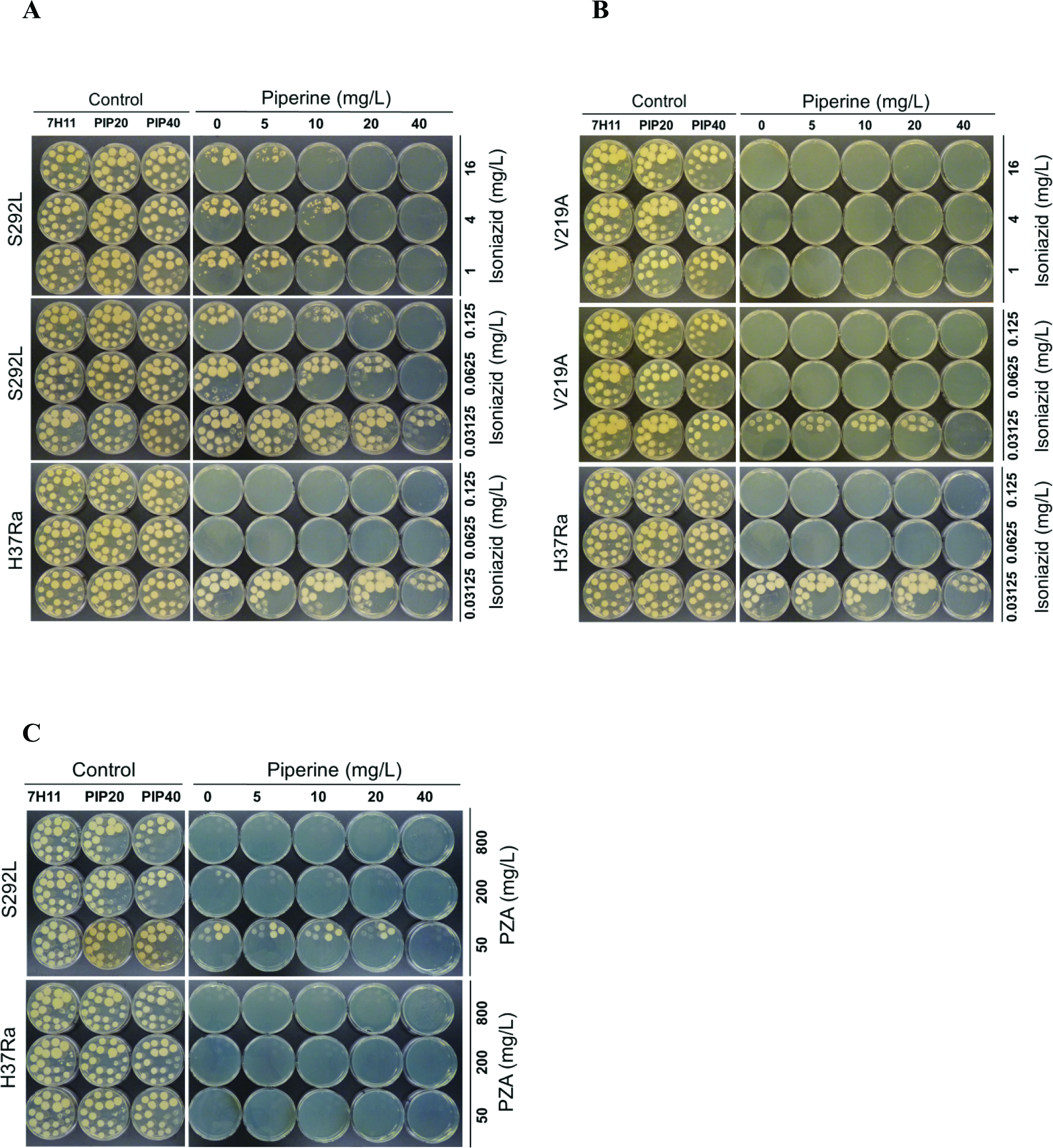
Piperine inhibited the INH (A) and PZA resistance (C) resistance in the Rv1258c S292L mutant, but not in the V219A mutant (B).

### PZA-resistant Rv1258c S292L mutant accumulates less drug POA

It has been shown that [^14^C]PZA is converted to [^14^C]POA in *M. tuberculosis* and accumulates in the cell to some extent due to a weak efflux of POA (13). To determine if Rv1258c V219A and S292L mutants, which are confirmed to be PZA-resistant in susceptibility testing (Fig. 3), can pump [^14^C]PZA out of the cell more effectively, we performed the [^14^C]PZA uptake experiment comparing the amount of [^14^C]POA accumulated in the cell in the Rv1258c mutants and that of the control parent strain. It was found that the intracellular concentration of POA in Rv1258c mutants was lower than that of PZA-susceptible *M. tuberculosis* H37Ra (Fig. 6), indicating that PZA accumulated to a less extent in the PZA-resistant Rv1258c S292L and V219A mutants, which is an indication of higher efflux activity of the Rv1258c mutants than the parent strain. In addition, the extent of [^14^C]POA accumulation was in accordance with the degree of resistance of the Rv1258c mutants, with the S292L mutant accumulating less POA than the V219A mutant, while both mutants accumulated less POA than the parent strain (Fig. 6B).

**Figure 6.**
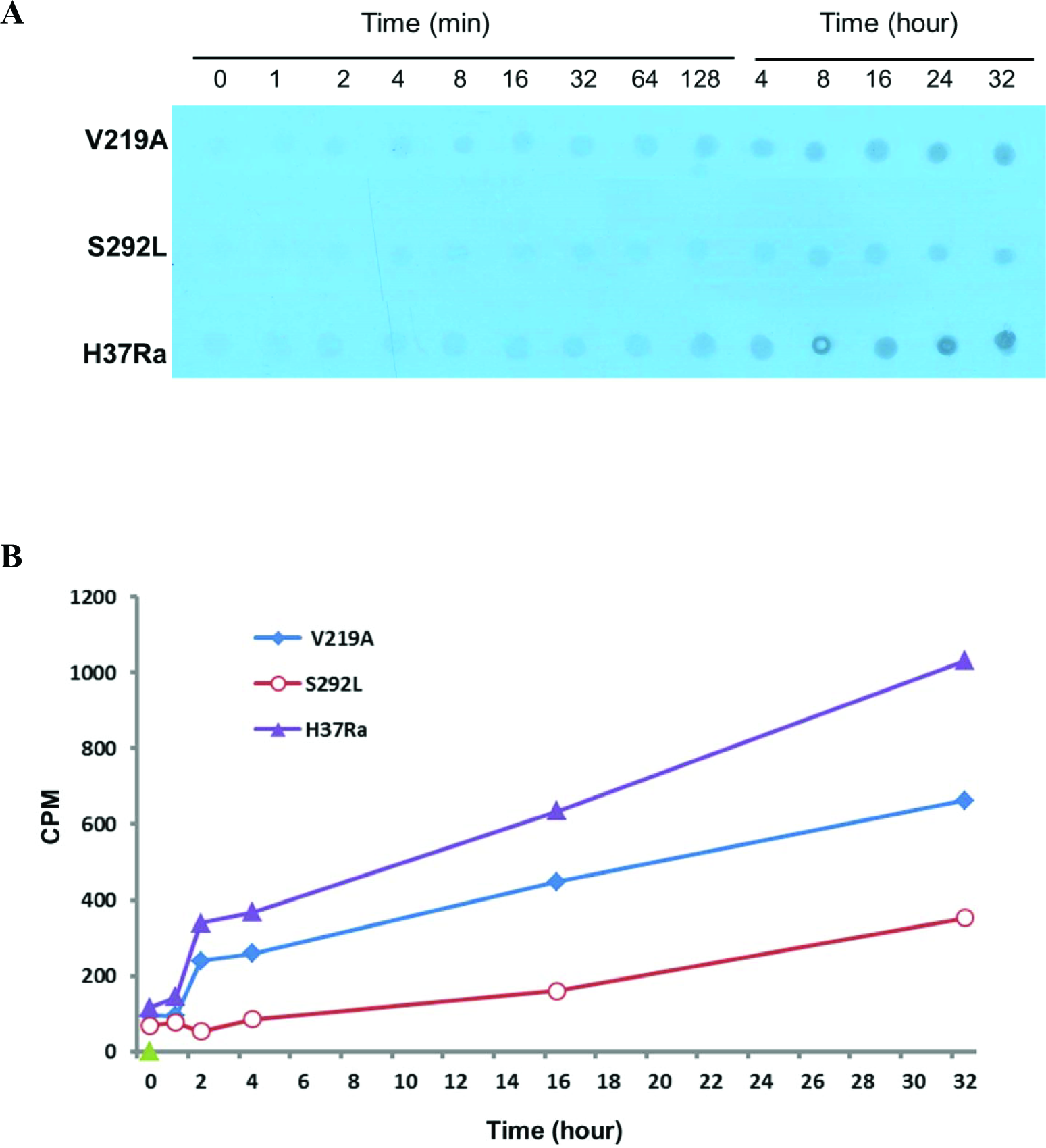
Comparison of PZA/POA accumulation in Rv1258c mutants. [^14^C]PZA was added to 5×10^9^ *M. tuberculosis* H37Ra at a concentration of 1μCi/ml at pH 6.6. At different times after PZA addition, portions of bacterial cells were removed and washed by filtration using phosphate buffered saline (pH 6.6) as described in Methods. The results of PZA/POA accumulation are shown by autoradiography (A) and scintillation counting (B). The S292L mutant accumulated less drug than the V219A mutant, while both mutants accumulated less drug than the parent strain *M. tuberculosis* H37Ra.

In conclusion, we demonstrate that point mutations in efflux pump Rv1258c found in clinical isolates can play an important role in conferring clinically relevant drug resistance to multiple drugs including PZA, SM and INH. Our findings could explain some previously unaccounted drug resistance in drug-resistant clinical strains and indicate efflux pump mutations may need to be taken into consideration for improved molecular detection of drug resistance in *M. tuberculosis*. Furthermore, efflux pump inhibitor piperine may be used as adjunct for possible more effective treatment of multi-drug resistant *M. tuberculosis* in future studies.

## FUNDING

This work was supported by the US-Russia (NIH-RFBR) Collaborative Research Partnership on the Prevention and Treatment of HIV/AIDS and Comorbidities grants from NIH AI108535 and AI099512, and the Russian Foundation for Basic Research (RFBR) grant 13-04-91444.

